# The Effect of Berberine on the Transcriptome and Proteome of *E. coli*

**DOI:** 10.1101/318733

**Authors:** Zhang Shilei, Jia Ze, Zhai Xianghe, Wang Chunguang, Zhang Tie

## Abstract

Berberine is commonly used to treat diarrhea in China, and the antibacterial properties of berberine have been confirmed. High-throughput sequencing technology was used to explore the changes induced in *E. coli* by berberine. After treatment with berberine, the expression of RstA and YbjG were found to be significantly different by RNA-seq and quantitative proteomics. However, the levels of MdtA, PmrA, LolD, LptG, MlaB, RcsF, and DppB were found to be significantly different by quantitative proteomics. Transcriptome sequencing did not yield as many results as proteome sequencing. The results of small RNA prediction showed increased sRNA00002 levels. The study showed that the differentially expressed mRNAs and proteins were associated with multidrug-resistant efflux systems. It can be inferred that berberine reduces *E. coli* antibiotic resistance. The results of this study are undoubtedly valuable to other researchers.

## 1 Introduction

Berberine, an alkaloid originally extracted from *Coptis chinensis* and other herb plants, significantly inhibits the growth of bacteria, viruses and protozoa^1,2^. Researchers have found that enhancement of the fluconazole-susceptibility of *Candida tropicalis* is mediated by berberine ^3^. Wang Jie et al. determined the minimum inhibitory concentrations (MIC) of *Belamcanda chinensis*, *C. chinensis* Franch, and the Sanhuang decoction (*Scutellaria baicalensis* Georgi, *C. chinensis* Franch, *Rheum palmatum* L.) against multidrug-resistant *Acinetobacter baumannii*, and the results demonstrated that the reversal rates were 4.2%, 2.8% and 11.1%, respectively ^4^. Eumkeb G. et al. claimed that galangin, quercetin and baicalein all had the potential to eliminate the tolerance of penicillin-resistant *Staphylococcus aureus* to AMP antibiotics ^5^. Bouhdid S. et al. discovered that thymol and carvacrol in oregano are involved in the regulation of membrane permeability in bacteria and function by destroying the cell membrane structure and increasing sensitivity of multidrug-resistant bacteria to antibiotics ^6^. Other studies have proven that baicalin can inhibit bacterial membrane systems ^7^. When penicillin-resistant *S. aureus* was treated with chlorogenic acid, the cell membrane composition of the bacteria changed, and the susceptibility of *S. aureus* to penicillin improved ^8^. Modern pharmacologists have shown that bacteria become sensitive to a variety of antibiotics when treated with berberine. Berberine can inhibit the function of bacterial cell membranes ^9^, reduce the lipopolysaccharide content of bacterial cell walls, and increase the susceptibility of bacteria to antibiotics ^10,11^. In addition, the glycolic acid oxidation and pyruvate oxidation pathways are inhibited by berberine during the process of glucose metabolism 12. However, the mechanism via which berberine mediates the enhancement of the antibiotic-susceptibility of *Escherichia coli* remains unclear.

High-throughput sequencing technology is widely used in scientific research. However, there remains a gap between traditional Chinese medicinal research and high-throughput sequencing technology. There has been no research to identify the role of berberine in *E. coli* via transcriptome and proteome analyses. To explore the changes mediated by berberine in *E. coli*, RNA-seq and quantitative proteomic techniques were used in this study. We were able to obtain complete transcriptomic and proteomic data for *E. coli* with or without berberine treatment. In this study, we found that the expression of *RstA* (b1608) and *YbjG* (b0841) was significantly different by RNA-seq and quantitative proteomic analyses. However, the expression of *MdtA* (b2074), *PmrA* (b4113), *LolD* (b1117), *LptG* (b4262), *MlaB* (b3191), *RcsF* (b0196), and *DppB* (b3543) was significantly different by quantitative proteomic analysis. The results of small RNA prediction showed that the level of sRNA00002 increases. Gene Ontology (GO) and Kyoto Encyclopedia of Genes and Genomes (KEGG) enrichment analyses showed that these significantly differentially expressed genes (DEGs) were primarily annotated as being involved in cell surface signaling, ion binding, and lipid metabolism, while the differentially expresses proteins were annotated as membrane proteins, cell wall synthesis proteins and cell membrane synthesis proteins.

## 2 Materials and Methods

A strain of E. coli used in this study was isolated from the Animal Hospital of Hebei Agricultural University, Baoding.

### 2.1 RNA-seq

RNA degradation and contamination were monitored by agarose gel electrophoresis, and purity was checked by using a Nano Photometer spectrophotometer (Nano drop 2000, Roche, Switzerland), while concentration was measured by using a Qubit RNA Assay Kit (Thermo, US) with a Qubit 2.0 fluorometer. Then, RNA integrity (Life Technologies, CA) was assessed by an Agilent Bioanalyzer 2100 system (Agilent, US). After passing the test, sequencing libraries were generated by the NEBNext Ultra Directional RNA Library Prep Kit (Biolabs, UK), after which the library fragments were purified with an AMPure XP system (Beckman, US). In addition, library quality was assessed on an Agilent Bioanalyzer 2100 system. After inspection, the library was pooled for Illumina HiSeqTM 2500 sequencing (Illumia, US).

#### 2.1.1 Mapping reads to the reference genome

Reference genome and gene model annotation files were downloaded directly from the genome website. Bowtie2-2.2.3 was used to both building an index of the reference genome and align clean reads to the reference genome. In this study, the reference genome selected for use was that of *E. coli* str. K-12 substr. mg1655 (ftp://ftp.ensemblgenomes.org/pub/bacteria/release31/fasta/bacteria_0_collection/escherichia_coli_str_k_12_substr_mg1655/dna/).

#### 2.1.2 Small RNA prediction and changes in expression

Non-coding RNAs with lengths of 50 to 500 nt are generally defined as small RNA. New intergenic region transcripts were found by using Rockhopper software. The newly predicted transcribed region was annotated by the BLASTx and the non-redundant (NR) nucleotide library. Non-commented transcripts were non-coding for small RNA. Secondary structure prediction and target gene prediction of the candidate small RNA were carried out by RNAfold software and IntaRNA software, respectively.

#### 2.1.3 Quantitative analysis of gene expression levels

Fragments per kilobase of exon per million fragments mapped (FPKM), which considers the effect of sequencing depth and gene length for read counts at the same time, is currently the most commonly used method for estimating gene expression levels. HTSeq v0.6.1 was used to count the number of reads mapped to each gene. The FPKM of each gene was calculated based on the length of the gene and read counts mapped to the gene. Differentially expressed gene sequencing (DEGseq) analysis of the two conditions was performed by the DEGseq R package. The threshold for significantly different expression was a P-value less than 0.005 and 2-fold change (FC) in threshold conditions (log2(FC)>1).

#### 2.1.4 Gene Ontology (GO) and Kyoto Encyclopedia of Genes and Genomes (KEGG) enrichment analyses of differentially expressed genes (DEGs)

GO enrichment analysis of differentially expressed genes was implemented by the GOseq R package, in which gene length bias was corrected. GO terms with corrected P-values less than 0.05 were considered significantly enriched with differentially expressed genes.

KEGG is a database resource for understanding high-level functions and utilities of biological systems, such as cells, organisms and ecosystems, from molecular-level information, especially from large-scale molecular datasets generated by genome sequencing and other high-throughput experimental technologies (http://www.genome.jp/kegg/). We used KOBAS software to test the statistical enrichment of differentially expressed genes in KEGG pathways.

#### 2.1.5 Fluorescence-based real-time quantitative PCR (RT-qPCR) for validation of DEGs

To verify the reliability of RNA-seq results, 8 differentially expressed genes were selected for RT-qPCR validation (Table 1).

Table 1 Primer sequences used for RT-qPCR

### 2.2 Quantitative proteomic sequencing

The total proteins were analyzed by the Bradford protein assay and SDS-PAGE. Mass spectrometry was performed after passing the test assay. The proteins were subjected to capillary high-performance liquid chromatography (HPLC) on an Easy-nLC 1000 instrument after reductive alkylation and trypsin hydrolysis and then subjected to mass spectrometry using a Q Exactive HF mass spectrometer.

#### 2.2.1 Mass spectrometric analysis and identification of differentially expressed proteins

This study used the UniProt *Escherichia coli* database. The original data were analyzed by Mascot and Proteome Discoverer software, while Proteome Discover 2.0 software combined with Mascot’s algorithm was used for qualitative and quantitative calculation of proteomics data. The extracted proteins were analyzed by mass spectrometry, and the results were filtered with a peptide false discovery rate (FDR) less than 0.01. The threshold criterion for significant differences in the ratios of the protein levels identified was 1.5 FC (FC≥1.5 is upregulated; FC≤0.667 is downregulated; 0.667<FC<1.5 implies no significant change).

#### 2.2.2 GO and KEGG enrichment analyses of differentially expressed proteins

Differentially expressed proteins were analyzed by GO functional annotation analysis and KEGG pathway annotation analysis to identify significantly enriched GO functions and KEGG pathways. Terms with P-values less than 0.05 were considered significantly enriched with differentially expressed genes.

## 3 Results

### 3.1 Isolation, Identification and Screening of *E. coli*

Coli0 was identified as *E. coli*. To investigate the effect of berberine, *E. coli* was treated with berberine at 250 μg/mL (named Coli1). RNA and protein were extracted for RNA-seq and quantitative proteomic studies.

### 3.2 RNA-seq

To determine the effect of berberine on bacterial mRNA, we performed transcriptase sequencing. By RNA-seq, 14.51 million and 13.71 million clean reads were obtained for Coli0 and Coli1, respectively, and 92.18%, 93.97% were mapped to the reference genomes of Coli0 and Coli1, respectively. The Q20 and Q30 of Coli0 and Coli1 were 98.84%, 98.76% and 96.82%, 96.63%, respectively. There were 45 DEGs between Coli0 and Coli1. Among these DEGs, 30 genes were upregulated, and 15 genes were downregulated (Supporting Table 1, Fig 1).

**Fig 1.**
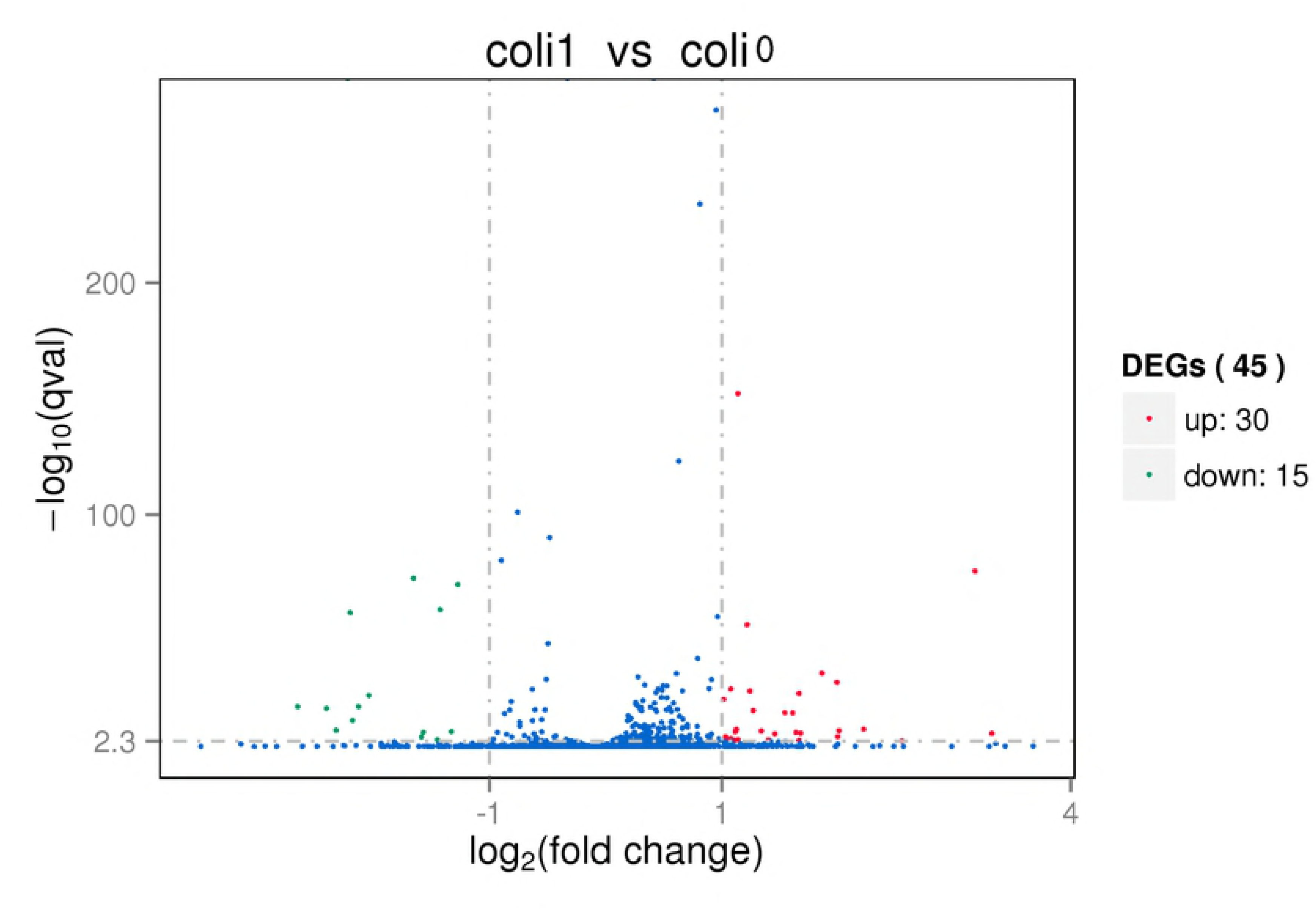
Volcano plot of DEGs between Coli1 and Coli0

#### 3.2.1 Small RNA prediction

New intergenic region transcripts were identified by comparison of the BLASTx and NR libraries. The newly predicted transcribed region was annotated, and the unlabeled transcripts were used as candidate non-coding small RNAs. Secondary structure prediction and target gene prediction of the candidate small RNAs were carried out by RNAfold software and IntaRNA software, respectively. A 72-bp small RNA was predicted, namely, sRNA00002 (Fig 2). The target gene of sRNA00002 is *DppB*; the complementary position is 849-903; and the dissociation energy is -13.3768 kcal/mol. In berberine-treated *E. coli*, the expression of sRNA00002 increased from 619.74 FPKM to 664.15 FPKM.

**Fig 2.**
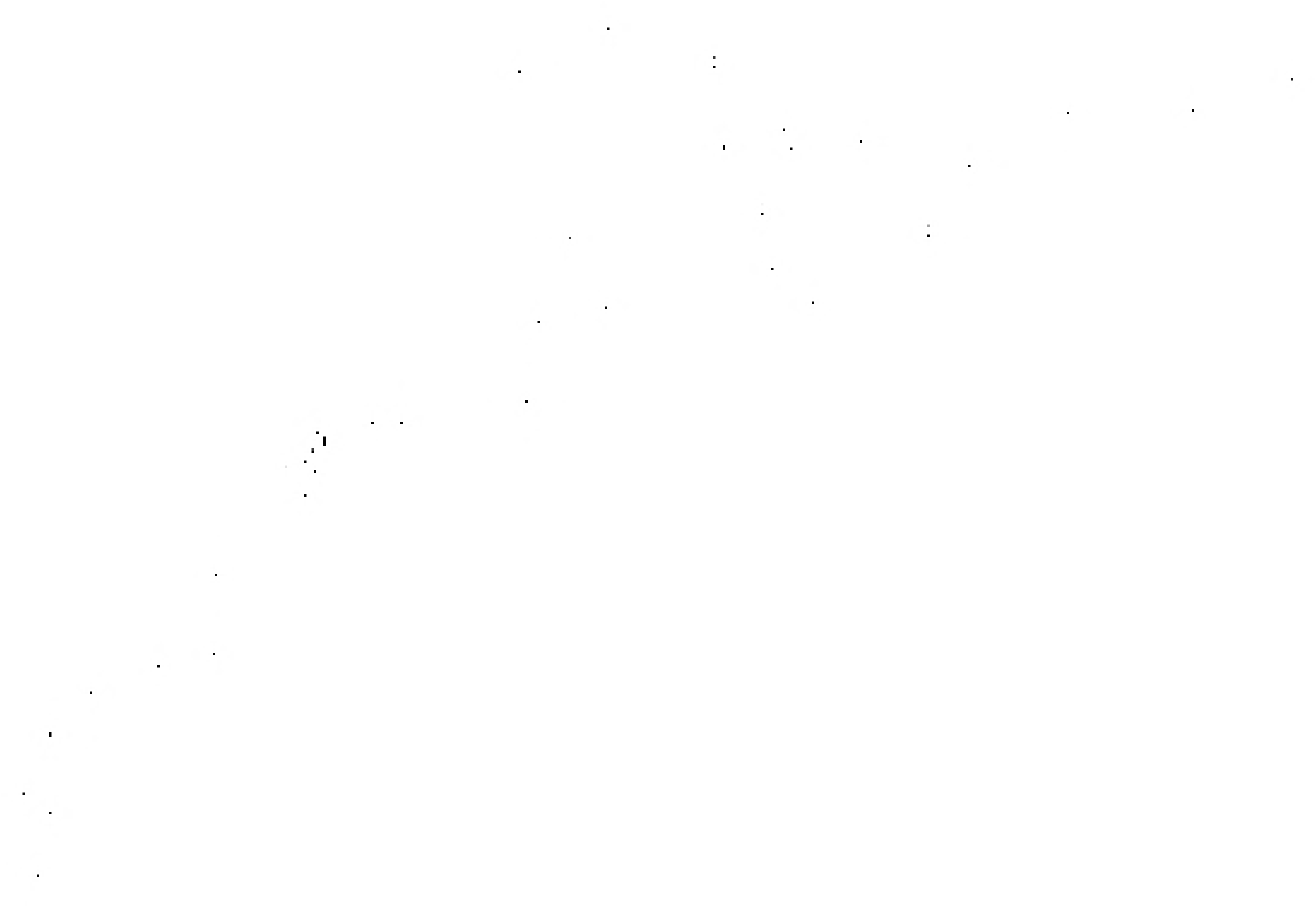
sRNA00002 secondary structure prediction

#### 3.2.2 Analysis of GO functions at the RNA level

To investigate the biological processes associated with berberine-induced levofloxacin resistance in *E. coli*, we performed GO enrichment analysis on DEGs detected by RNA-seq. The enriched GO functions included cell surface receptor signaling pathway, lipid modification, positive regulation of immune system process, antigen receptor-mediated signaling pathway, regulation of antigen receptor-mediated signaling pathway, ion binding, Wnt receptor signaling pathway, cellular lipid metabolic process, lipid phosphorylation, regulation of signal transduction, regulation of cell communication, regulation of signaling, single-organism metabolic process, DNA polymerase III complex, DNA polymerase complex, lipid metabolic process, signal transduction, anion binding, signaling, and single-organism signaling (Table 2).

Table 2 GO function enrichment at the RNA level in Coli0 and Coli1

#### 3.2.3 Analysis of KEGG pathways at the RNA level

To study the metabolic pathways associated with levofloxacin resistance in *E. coli*, KOBAS software was utilized to detect metabolic pathways affected by the DEGs. Seven significant KEGG pathways were identified as being enriched, namely, fatty acid degradation, glutathione metabolism, fatty acid metabolism, peptidoglycan biosynthesis, microbial metabolism in diverse environments, metabolic pathways, and two-component system (Table 3).

Table 3 KEGG enrichment at the RNA level in Coli0 and Coli1

#### 3.2.4 RT-qPCR

Eight genes were randomly selected from 45 significantly differentially expressed genes for validation. Comparison of the sequencing results with the multiple expression values of each gene obtained by RT-qPCR demonstrated that the trend of gene expression remains roughly the same; however, there is a difference between the two trends. This discrepancy may be a result of the differences in instrumentation, reagents, operating errors, or algorithms (Fig 3).

**Fig 3.**
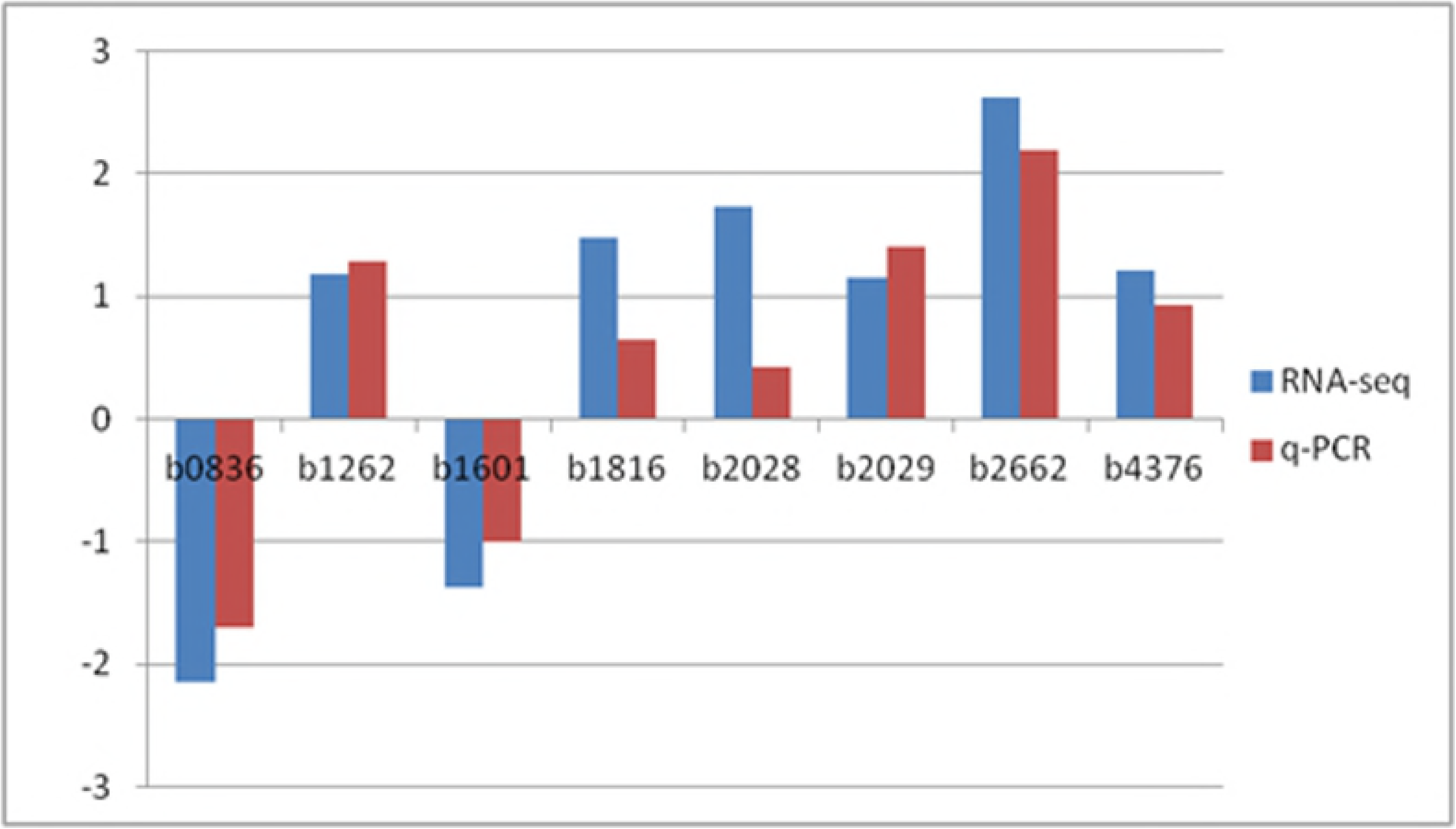
Expression levels of eight genes from the Coli0 and Coli1 determined by RT-qPCR and RNA-seq. Positive values on the Y axis represent upregulation, and negative values represent downregulation.

### 3.3 Quantitative proteomic sequencing

To characterize the effect of berberine on bacterial protein synthesis, we used label-free proteomic sequencing analysis. A total of 2003 proteins were obtained from quantitative proteomic sequencing. There were 136 unique proteins in Coli0, 162 unique proteins in Coli1. There were 331 downregulated proteins and 291 upregulated proteins in Coli1. To study the changes in resistance-related proteins in thee mutant *E. coli*, we analyzed the differential expression of membrane proteins, cell wall synthesis proteins and cell membrane synthesis proteins.

#### 3.3.1 GO Analysis of antibiotic-resistant proteins

In the GO function enrichment analysis, 37 terms were significantly enriched, including plasma membrane, response to metal ion, response to inorganic substance, metal ion transport, glycerol catabolic process, organic acid transmembrane transporter activity, carboxylic acid transmembrane transporter activity, ion transport, nitrogen compound transport, integral component of membrane, transporter activity, organic anion transmembrane transporter activity, anion transport, organic anion transport, alditol metabolic process, carboxylic acid transport, glycerol metabolic process, cell periphery, substrate-specific transmembrane transporter activity, transmembrane transporter activity, intrinsic component of membrane, substrate-specific transporter activity, alditol catabolic process, cellular response to inorganic substance, transcription factor activity, sequence-specific DNA binding, nucleic acid binding transcription factor activity, plasma membrane part, secondary active transmembrane transporter activity, transport, membrane part, response to transition metal nanoparticle, phenol-containing compound metabolic process, cell projection organization, racemase and epimerase activity, acting on amino acids and derivatives, metal ion transmembrane transporter activity, amino acid transport, and heme binding (Table 4).

Table 4 GO function enrichment at the protein level in Coli0 and Coli1

#### 3.3.2 KEGG analysis of antibiotic-resistant proteins

The enriched KEGG pathways included ABC transporters; alanine, aspartate and glutamate metabolism; quorum sensing; glycerophospholipid metabolism; butanoate metabolism; nitrotoluene degradation; beta-alanine metabolism; two-component system; glycerolipid metabolism; peptidoglycan biosynthesis; cationic antimicrobial peptide (CAMP) resistance; beta-lactam resistance; vancomycin resistance; lipopolysaccharide biosynthesis; metabolism of xenobiotics by cytochrome P450; drug metabolism – cytochrome P450; streptomycin biosynthesis; bacterial secretion system (Table 5>).

Table 5 KEGG enrichment at the protein level in Coli0 and Coli1

## 4 Discussion

The response regulator of *RstAB* (*RstA*) was significantly downregulated. The *RstAB* system is a bacterial iron uptake system, and downregulation of this system may lead to inhibition of bacterial iron absorption. The *RstAB* system is associated with assistance to harsh environments^13^. However, the relationship between *RstAB* and antibiotic resistance in bacteria has not been proven. Considering the use of berberine, the significant reduction in *RstA* expression might be associated with the stimulation of berberine.

Activation of the capsule synthesis promoter (*RcsF*) was significantly downregulated. *RcsF* is involved in cell wall synthesis, and downregulation of *RcsF* leads to the inhibition of cell wall synthesis. Modification of the *RcsF* protein may act as an activating signal, so the expression of *RcsF* was not significantly different when measured by RNA-seq^14,15^. Undecaprenyl-diphosphatase (*YbjG*) was significantly downregulated. *YbjG* is essential for the synthesis of undecaprenyl phosphate, a C55 lipid carrier for cell wall synthesis^16^. In addition, *YbjG* is similar to the bacitracin-resistance protein *BcrC* of *Bacillus licheniformis*^17^. Touzé et al. inhibited the phosphorylation of lipid A in bacterial cell walls by using bacitracin, suggesting that lipid A phosphorylation in bacterial cell walls is directly associated with undecaprenyl pyrophosphate synthesis^18^. Incomplete cell walls weaken the antibiotic resistance of *E. coli*. This study proved that berberine can significantly inhibit bacterial cell wall synthesis.

The multidrug resistance protein (*MdtA*) was significantly downregulated. *MdtA* can cause bacteria to become dormant and resume growth after the antibiotic is no longer present. Bacteria can escape killing by antibiotics via transient dormancy, which occurs only during bacterial proliferation. The principle behind this phenomenon is that in the presence of antibiotics, the bacterial cells themselves produces a toxin protein that allows the bacteria to enter a dormant state from a growing state to achieve the goal of resisting antibiotics^19^.

The transcriptional regulator of the multidrug resistance efflux pump *PmrAB* (*PmrA*) was significantly downregulated. *PmrAB* is the regulator of lipopolysaccharide (LPS) modification in *E. coli,* and *PmrA* is the transcriptional regulator of this system^20^. The expression of *PmrAB* reduces the negative charge on the bacterial outer membrane, which affects the binding of bacteria to cationic antibiotics, leading to drug resistance^21^. Arroyo L. A. found that the change in *PmrAB* expression was associated with increased polymyxin resistance^22^. Interestingly, a decrease in *PmrA* expression attenuates the efflux of all drugs from *E. coli*, not only levofloxacin.

In this experiment, the expression levels of the ATP-binding protein of the lipoprotein release system (*LolD*) and permeation-associated enzyme of the lipopolysaccharide output system (*LptG*) were significantly decreased and that of the binding protein of phospholipid ABC transporter (*MlaB*) was significantly decreased. *LolD* provides energy for the transport of lipoproteins, and the absence of *LolD* will decrease the rate of transport of lipoproteins to the outer membrane^23^. *Mla* is involved in maintaining outer membrane lipid asymmetry via the transport of aberrantly localized phospholipids from the outer membrane to the inner membrane, while *MlaB* plays critical roles in both the assembly and activity of the transporter^24^. In gram-negative bacteria, lipid asymmetry is critical to the function of the outer membrane as a selectively permeable barrier. LPS can act against many antimicrobials. To assemble LPS on their surfaces, bacteria must extract LPS from the inner membrane to the outer membrane. The *Lpt* system transports LPS by using the ATPase *LptB* and the transmembrane domains *LptFG*^25^. Decreased *LloD*, *LptG* and *MlaB* expression will destroy cell membrane stability and weaken the role of the cell membrane barrier.

The expression of dipeptide ABC transporter permease (*DppB*) was significantly reduced. Peptide transporters play important roles in bacterial growth, not only in the supply of nutrients to bacteria but also in many processes of the bacterial lifecycle, such as signal transduction and chemotaxis^26,27^. There have been no studies to prove that the peptide transport system is associated with antibiotic resistance. The small RNA that regulates *DppB* was also identified for the first time. We have not yet studied the peptide transport system further, but this system is a promising candidate as a system that is involved in *E. coli* antibiotic resistance. The newly discovered small RNA might be able to explain this phenomenon.

## Supplementary Material

Supplementary Table Differentially expressed genes from Coli0 and Coli1. Between Coli0 and Coli1, the two *E. coli* cultures, a total of 45 genes were differentially expressed; “readcount” represents the relative expression level (FPKM); “log2 Fold change” represents the multiple of differential expression; negative values represent downregulates genes; positive values represent upregulated genes; “pvalue” and “qvalue” less than 0.01 represent significant differences.

